# Contrasting effects of acute and chronic stress on the transcriptome, epigenome, and immune response of Atlantic salmon

**DOI:** 10.1101/319285

**Authors:** Tamsyn M. Uren Webster, Deiene Rodriguez-Barreto, Samuel A.M. Martin, Cock van Oosterhout, Pablo Orozco-terWengel, Joanne Cable, Alastair Hamilton, Carlos Garcia de Leaniz, Sofia Consuegra

## Abstract

Early-life stress can have long-lasting effects on immunity, but the underlying molecular mechanisms are unclear. We examined the effects of acute stress (cold-shock during embryogenesis) and chronic stress (absence of tank enrichment during larval-stage) on the gill transcriptome and methylome of Atlantic salmon four months after hatching. While only chronic stress induced pronounced transcriptional effects, both acute and chronic stress caused lasting, and contrasting, changes in the methylome. Crucially, we found that acute stress enhanced immune response to a pathogenic challenge (lipopolysaccharide), while chronic stress suppressed it. We identified stress-induced changes in promoter or gene-body methylation that were associated with altered expression for a small proportion of genes, and also evidence of wider epigenetic regulation within signalling pathways involved in immune response. Our study suggests that early-life stress can affect immuno-competence through epigenetic mechanisms, a finding that could open the way for improved stress and disease management of farmed fish.

## Introduction

Stress response is a fundamental survival mechanism that provides a critical adaptive response to many environmental challenges, but may also compromise the immune system (Calcagni & Elenkov 2006; Chrousos 2009). The precise impacts of environmental stress on immune function are highly dependent on the timing, duration, magnitude and nature of the stressor and stress response (Tort 2011). Chronic stressors, lasting for days or weeks, can dysregulate immune response by inducing long-term changes in energetic metabolism, persistent low level inflammation, and by suppressing the release of immune cells and cytokines (Dhabhar 2014). In contrast, acute stressors, lasting for minutes to hours, are less likely to impair immune function, and may even enhance immune response by stimulating the maturation, secretion and redistribution of immune cells and cytokines (Dhabhar 2009).

For vertebrates, early life stages may be particularly sensitive to environmental stress, due to developmental plasticity during critical periods of differentiation and maturation of the nervous and immune systems (Cao-Lei et al. 2017). In mammalian systems, it is well established that early-life stress can have long-lasting adverse effects on health and disease susceptibility. For example, maternal stress during pregnancy predisposes the offspring to developmental, immunological and behavioural abnormalities throughout their life, and post-natal trauma is associated with an increased risk of depression, obesity, diabetes and cardiovascular disease (Silberman et al. 2016; Vaiserman 2015). However, exposure to mild stress during early life may have beneficial effects later in life, a phenomenon known as hormesis (Mattson & Calabrese 2009). The molecular mechanisms underlying this effect are not fully understood, but hormesis can enhance immune function as part of a primed, more efficient response to future stressors (Monaghan & Haussmann 2015), and could even be harnessed in a clinical setting to boost protective immunity (Dhabhar 2014).

Environmental stress also induces changes in the epigenome, providing a mechanism by which stress can have long-term effects on transcriptional regulation and phenotype throughout an individual’s lifetime and, in some cases, on its progeny (Cao-Lei et al. 2017; Silberman et al. 2016). Epigenetic modifications following exposure to stress during early life are known to induce lasting transcriptional and structural changes in the mammalian brain (Cruceanu et al. 2017; Hunter et al. 2015; Nasca et al. 2015a; Nasca et al. 2015b). On a gene-specific basis, promoter silencing activity, whereby DNA methylation in promoter regions negatively regulates gene transcription, has been demonstrated to mediate lasting effects of stress on physiology, behaviour and psychiatric disorders (Cruceanu et al. 2017; McGowan et al. 2009; Nasca et al. 2015b). However, at the genome-wide level, the association between DNA methylation and gene expression is not straightforward. Complex interactions between different targets, cell types and layers of epigenetic regulation may facilitate wide, indirect effects of environmental stress (Hunter et al. 2015; Silberman et al. 2016). Beyond these critical effects on brain and behaviour, stress is also likely to have far-reaching effects on whole-organism physiology, including immunity, metabolism, nutrition and reproduction, which have been largely unexplored (Papadopoulou et al. 2015).

Fish are subjected to high levels of stress in aquaculture systems due to confinement, handling and environmental mismatch, which can impair immuno-competence and increase disease susceptibility (Vindas et al. 2016). Improving stress and disease resistance is a critical priority for the sustainable growth of aquaculture, which needs to provide a reliable and safe source of food for a growing human population and reduce impacts on welfare and the environment (Stentiford et al. 2017). Stress has well known effects on fish (e.g. Schreck et al. 2016; Tort 2011), but little is known about how stress experienced during early development can affect later life health outcomes, or the underlying molecular mechanisms responsible. During early life, many fish species experience critical periods for survival that coincide with the transition from endogenous to exogenous feeding and development of the immune system, when they are especially sensitive to stress (Castro et al. 2015; Elliott 1989). Recent research also suggests that stress modifies the fish epigenome in a developmental-stage specific manner, with these early life stages displaying a heightened period of epigenetic sensitivity (Moghadam et al. 2017). Therefore, we hypothesised that chronic stress experienced during early development would adversely affect immune function, while short-lived, mild stress could enhance immuno-competence, and that these effects might be mediated by epigenetic mechanisms. In order to test these hypotheses, we compared the effects of acute stress (cold shock and air exposure during late embryogenesis) and chronic stress (lack of tank enrichment during larval stage) on the gill transcriptome and methylome of Atlantic salmon (*Salmo salar*) fry, and also examined transcriptional immune response to a model pathogenic challenge (bacterial lipopolysaccharide, LPS).

## Results

### Survival and growth

Overall hatching success was 95.3 ± 1.1% and larval survival until 110 days post-hatching was 89.1 ± 1.1%, with no significant difference between the control, acute stress or chronic stress groups (hatching; *F*_*3,2*_*=1.38, P=0.377,* survival; *F*_*3,2*_*=0.19, P=0.836*). There was a significant effect of treatment on growth rate during pre- and early-feeding (748 and 1019 degree days (DD)), whereby fish exposed to chronic stress had significantly reduced mass compared to those in the control and acute stress groups, but this effect was no longer apparent at later sampling points (492 DD: *F*_*3,114*_*=0.37, P=0.718;* 748 DD: *F*_*3,114*_*=15.82, P=0.025;* 1019 DD: *F*_*3,114*_*=15.42, P=0.026;* 1323 DD: *F*_*3,114*_*=1.63, P=0.330,* 1532 DD: *F*_*3,114*_*=4.78, P=0.117;* Table S1), and there was no significant difference in condition factor between groups at the final sampling point *(F*_*3,114*_*=2.183, P=0.260*). Exposure to LPS for 24h caused no mortalities, or any apparent physiological or behavioural changes.

### Transcriptomic analysis

After quality filtering, an average of 27.8 million paired end RNA-seq reads (91.8%) were retained per sample. Of these, a total of 94.5% were mapped to the Atlantic salmon genome, including 84.1% unique alignments (Table S2). Following transcript reconstruction and novel assembly, we obtained a total of 201,433 transcripts in 104,528 putative loci, and statistical expression analysis was performed for 78,229 genes with nonzero read counts.

Expression analysis with DeSeq2 identified a total of 19 genes significantly differentially expressed (FDR <0.05) between the acute stress group and the control fish. In the chronically-stressed group, there were 206 differentially expressed genes compared to the control group, the vast majority (190, 92.2%) of which were up-regulated, and a functional analysis of these genes revealed strong enrichment of ribosome structural constituents and translation, as well as muscle development, energy metabolism and bacterial defence response (Figure S1).

LPS exposure had a very marked effect on gene transcription in the gills. MDS analysis based on the whole transcriptome clearly separated all LPS-exposed individuals from non-exposed fish (Figure 1a). The main effect of LPS exposure in the (non-stressed) control group consisted of 14,833 up-regulated and 10,636 down-regulated genes (FDR <0.05) in fish exposed to 20 μg/ml LPS compared to non-exposed fish (Figure S2). These included a large number of genes encoding proteins typically associated with inflammatory immune response including a large number of mucins; interleukins, interferons, TNF, chemokines and their regulatory factors and receptors; complement factors, immunoglobins and damage-inducible molecular chaperones. Fold changes for a selected list of these significantly differentially expressed genes with direct immune function is provided in the supporting information (Table S3). Overall functional analysis for the up-regulated genes revealed strong enrichment of GO terms related to cellular stress response (Figure S3). Enriched terms included those related to cell adhesion, cellular signal transduction, regulation of transcription, protein modification, response to LPS and response to cytokines (Biological Process), extracellular matrix (Cellular Component), and transcription factor activity, protein kinase/phosphatase activity, and binding of signalling molecules (Molecular Function). Enriched KEGG pathways included extracellular matrix-receptor interaction, as well as specific pathogen recognition pathways (NOD-like, RIG-like, TOLL-like receptor signalling). For down-regulated genes enriched terms were related to general cell maintenance processes. These included terms related to DNA-replication and repair, redox reactions, cell cycle and translation (Biological Process), ribosome and mitochondria (Cellular Component), ribosome constituent, redox activity and metabolic enzyme activity (Molecular Function), and DNA replication, ribosome and metabolic pathways (KEGG pathway).

**Figure 1.**
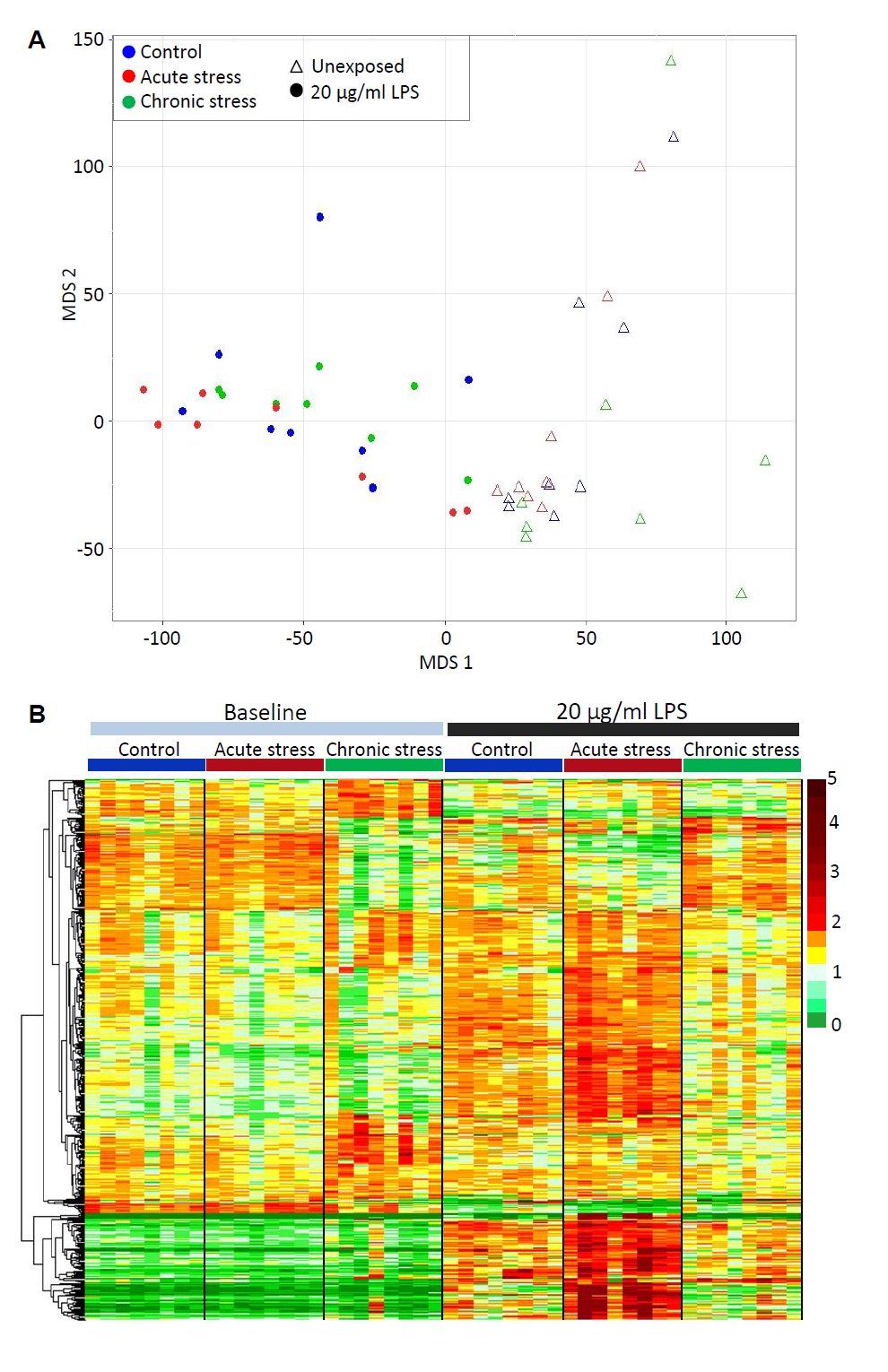
The impact of early life stress on transcriptional response to LPS. **A)** Multidimensional scaling analysis illustrating the very significant effect of exposure to 20 μg/ml LPS on the entire gill transcriptome (78,229 putative loci) of fish from all stress treatment groups. **B)** Heat map illustrating the expression of all genes for which a significant interaction between stress and LPS response was identified (516 genes). Data presented are read counts for each individual normalised by library size, and by mean expression for each gene. Hierarchical clustering was performed using an Euclidian distance metric.

We also identified a significant interaction between stress and transcriptional response to pathogen challenge. Compared to the main effect of LPS exposure identified in control fish (described above), we found that acute and chronic stress during early life altered transcriptional response to LPS in contrasting ways (Figure 1b). In the acutely stressed fish, a total of 194 genes were regulated by LPS in a significantly different way (FDR <0.05) compared to in control fish. Of these genes, the vast majority (140, 72.2%) were up-regulated relative to the control group in response to LPS and 41 (21.1%) were down-regulated relative to the control group in response to LPS. Only 9 (4.6%) genes that were regulated by LPS in the control group were not significantly responsive in the acutely-stressed group, and 4 (2.1%) genes were regulated in the opposite direction. Functional analysis of the genes with enhanced responsiveness to LPS in acutely-stressed fish revealed further enrichment of processes identified for the main LPS response, together with pathways related to lipid metabolism and mucin production (Figure S4a). For chronically-stressed fish, a total of 347 genes were expressed in a significantly different way following LPS exposure compared to the main effect of LPS exposure identified in non-stressed control fish. The majority of these genes (218, 62.8%) were not significantly regulated by LPS, or were significantly less responsive to LPS exposure relative to the control group (Figure 1b). Functional analysis of these less responsive genes revealed enrichment of a number of processes identified as part of the main LPS response (including cell adhesion, signal transduction, response to bacterium/lipopolysaccharide and p53 signalling), and amongst the more down-regulated genes there was a strong enrichment of ribosome and translation (Figure S4b). Only 28 (8.1%) and 57 (16.4%) genes, respectively, were up-regulated and down-regulated to a greater extent in chronically-stressed fish than in the control group, which is in stark contrast to the enhancive effect of acute stress on transcriptional response to LPS.

### Methylation analysis

After quality filtering, a total of 1534 million high quality single end RRBS reads, averaging 64 million/sample, were retained. A total of 90.6% of these were mapped to the reference genome, with a unique alignment rate of 43.5% (Table S4). Analysis of spike-in methylation controls revealed an overall bisulphite conversion efficiency of 99.7%, with 2.0% inappropriate conversion. In total, we identified 21.3 million CpG sites in our libraries, but only 1.1 million were covered in all samples, representing 2.75% of total CpGs in the Atlantic salmon genome, which is a comparable percentage to that previously reported for RRBS experiments in zebrafish (5.3%; Chatterjee et al. 2013) and rainbow trout (< 1%; Baerwald et al. 2016). Methylation analysis was conducted only using CpGs covered by at least 10 reads in all libraries (335,996 CpG sites).

The majority of the CpGs surveyed mapped to gene bodies (53%) or intergenic regions (45%), with 2% located in putative promoter regions. In terms of CpG context, 12% of CpGs were located in CpG islands, 17% in CpGshores, and 11% in CpGshelves. CpG methylation level dropped progressively in the region upstream of the TSS, then increased sharply within the gene body (Figure 2a). Genome-wide CpG methylation displayed a bimodal distribution, whereby the majority of CpGs within gene bodies (60.08%) were highly methylated (>80% methylation), while a large proportion (24.6%) were un-methylated or hypo-methylated (<20% methylation) (Figure 2b). Across the putative promoter regions, there was a higher proportion of hypo-methylated CpGs (64.1%) (Figure 2c). Genome-wide, the average methylation of all CpG sites was 74.87% ± 0.38 in the control group, 74.04% ± 0.71 in the acute stressed fish and 74.94% ± 0.82 in the chronically stressed group, with no significant difference among groups (*F*_*2,21*_*=0.55, P=0.59*).

**Figure 2.**
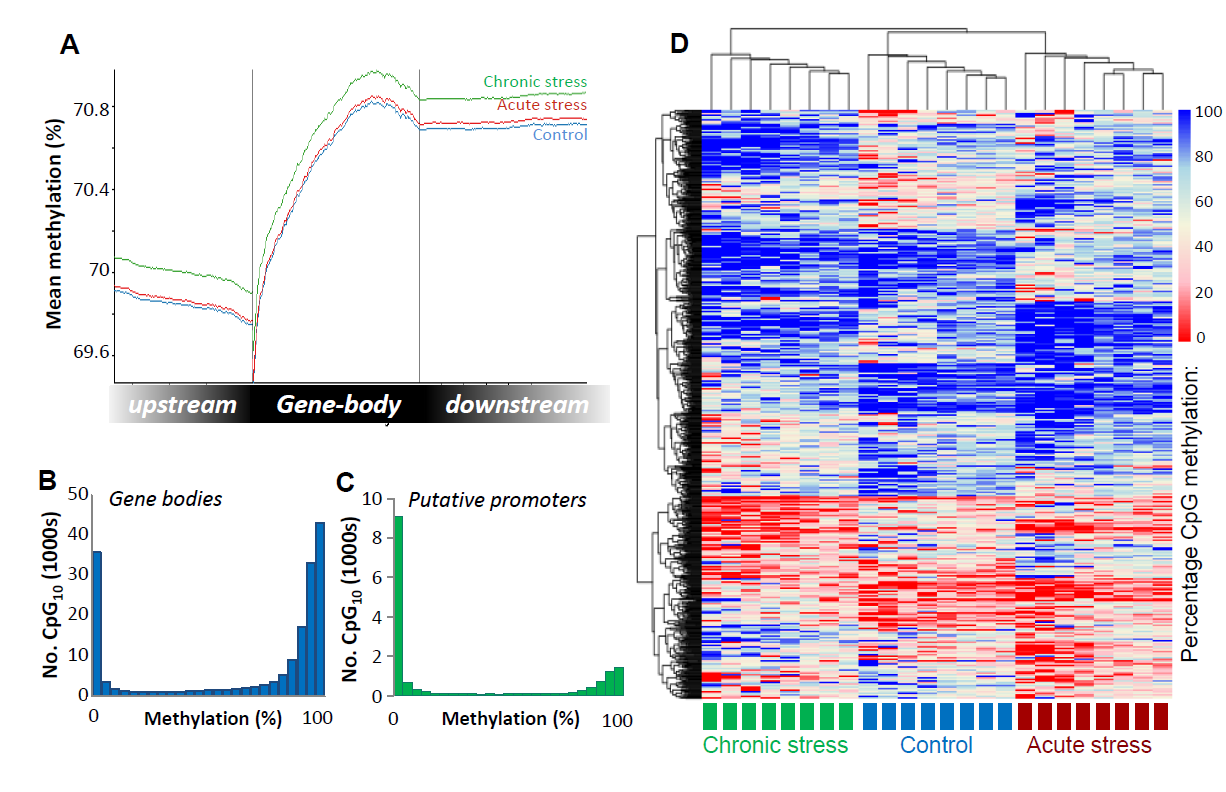
Visualisation of the Atlantic salmon gill methylome. **A)** Average CpG methylation percentage in gene bodies and within the 1.5 Kb upstream and downstream of the transcription start (TSS) and termination sites (TTS) for each stress group. **B-C)** Histograms of average methylation distribution within gene bodies and putative promoter regions. **D)** Heat map illustrating percentage methylation for all differentially methylated CpGs identified in each stress group (logistic regression q< 0.01 and |ΔM|>20%, and t.test p <0.01), using unsupervised hierarchical clustering.

Compared to the control fish, a total of 1895 and 1952 differentially methylated CpG sites (DMCpGs) were identified using logistic regression (FDR< 0.05, |ΔM|≥ 20%) in the acute stress and chronic stress groups, respectively. The genomic distribution and context of the DMCpGs largely mirrored the wider methylome landscape, with half of the DMCpGs overlapping intragenic regions (52%). DMCpGs overlapped or neighboured (up to 2 Kb from TSS) a total of 907 genes for the acute stress group, and 925 genes for the chronic stress group, including 242 common genes shared in both groups. For both stress groups, the most strongly enriched functional processes amongst these genes were related to cellular adhesion and cellular signalling pathways (Figure S5). These included terms related to cell adhesion, intracellular signal transduction, Rho protein signalling, calcium ion transport and signalling and ion transport (Biological Process), plasma membrane, cell junction, myosin complex (Cellular Component) and ion channel and transported activity, GTPase and guanyl-nucleotide exchange activity (Molecular Function). A more stringent list of 1004 total DMCpGs were identified across both types of stress using both statistical methods (logistic regression and t-tests). Unsupervised hierarchical clustering of these DMCpGs revealed a distinctive methylation profile for both the acutely and chronically stressed groups with respect to the controls and to each other, although there was greater resemblance between the control and acute stress groups (Figure 2d).

### Transcriptome-methylome integration

We examined the transcriptome wide association between gene expression and DNA methylation within putative promoter regions (p.promoters; windows from 1500bp upstream to 1000 bp downstream of the transcription start site (TSS)) and within gene bodies. There was a significant negative correlation between p.promoter mean methylation and gene expression (Spearman rho= −0.37; *P<*0.001), but no overall correlation between gene body methylation and gene expression (Spearman rho= −0.03; *P=*0.062). There was some evidence of a heterogeneous relationship between gene body methylation and gene expression. GAM analysis indicated that a small, but significant, part of gene expression was explained by the smooth component of gene body methylation (deviance explained = 1.93%, *P* < 0.001, Figure 3a,b).

**Figure 3.**
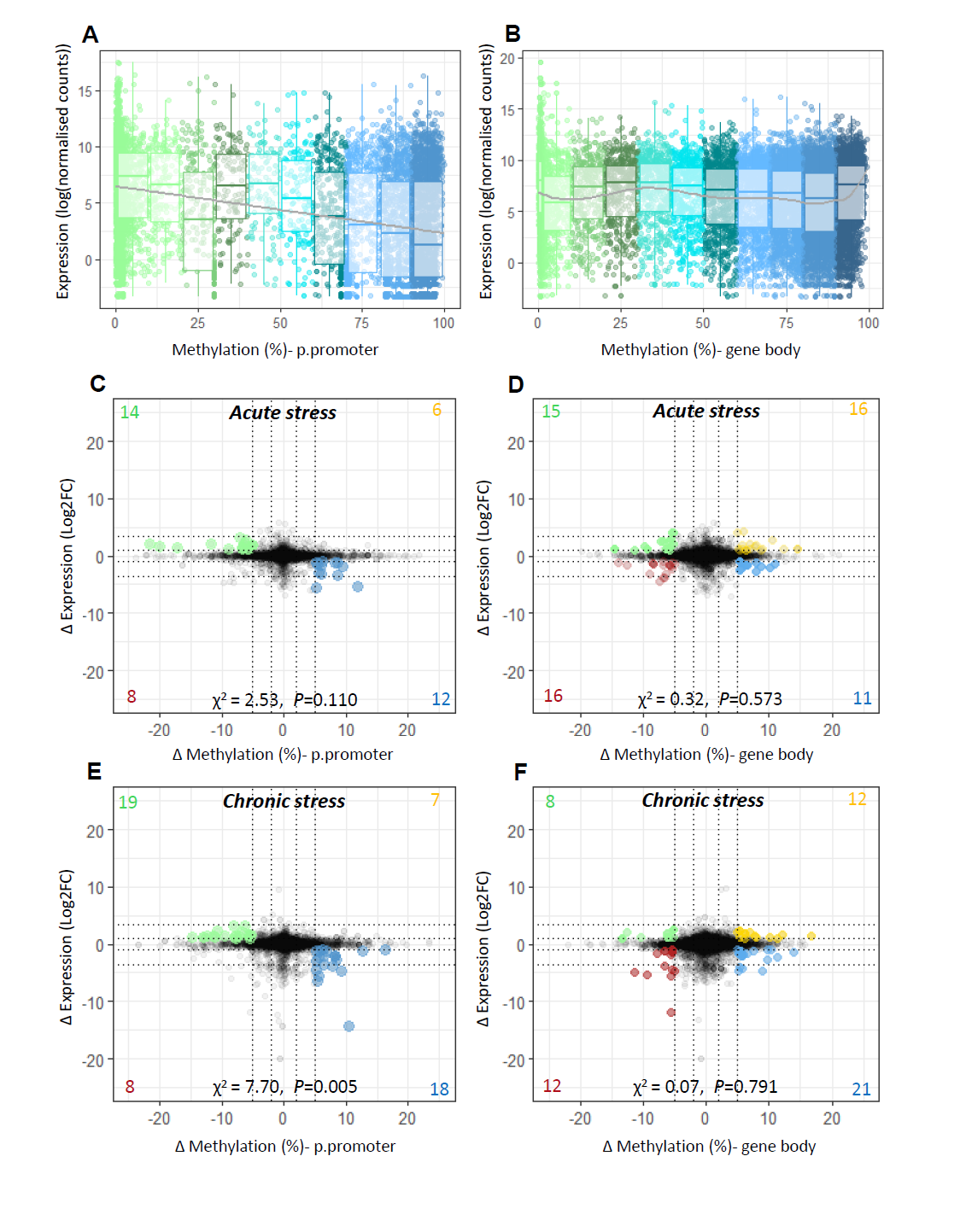
Integration of transcriptome and methylome. Scatterplot and boxplot displaying mean gene expression and mean DNA methylation for **A)** putative promoters and **B)** gene bodies in control fish. **C-F)** Starburst plots displaying the effect of stress on the transcriptome and the methylome. For each type of stress relative to the control group, change in gene expression (log2fold change) is plotted against change in DNA methylation (ΔM) for **(C;E)** *putative promoters* and **(D;F)** *gene bodies*. Highlighted dots denote genes with ΔM > 5% and |FC |> 2; a full list of highlighted genes is provided in Table S5-S6.

We then identified genes for which there was a notable effect of early life stress on both DNA methylation (p.promoters and gene bodies) and on gene expression (>2 FC delta expression and >5% methylation difference) at the baseline time-point (i.e. not exposed to LPS). For p.promoters, there was evidence of unequal distribution of genes between hyper-methylated/up-regulated, hypo-methylated/up-regulated, hyper-methylated/down-regulated and hypo-methylated/down-regulated groups (Acute stress: 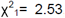, *P*=0.110, Chronic stress: 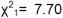, *P*=0.005) with greater numbers of genes with an inverse relationship between delta methylation and delta expression (Figure 3c,e; Table S5). Therefore, and given the overall negative relationship between methylation in p.promoter regions and gene expression we focused only on these genes with an inverse relationship. However, there was no evidence of a similar effect between delta gene body methylation and delta expression (Acute stress: 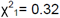, *P*=0.573, Chronic stress: 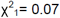, *P*=0.791), therefore we included all genes above the threshold (Figure 3d,f; Table S6). Combined functional analysis of these genes revealed enrichment of processes related to ion/calcium ion transport and signal transduction (Figure S6).

Finally, we examined the potential for stress-induced changes in baseline DNA methylation to contribute to the observed altered transcriptional response to LPS. We did not perform methylation analysis for the fish exposed to LPS, but we hypothesised that baseline promoter and/or gene body methylation status might influence the rapid transcriptional response induced by LPS exposure. Of the genes that showed a significant interaction between stress treatment and response to LPS, 28 (acute stress) and 57 (chronic stress) met the coverage criteria for targeted analysis of baseline p.promoter methylation (Figure S7). For acutely-stressed fish we identified three genes with hypo-methylation (|ΔM|>5%) and increased-expression in response to LPS treatment relative to the control group (*lrrn4cl, usp54a, st3gal1l3*), and for chronically-stressed fish we identified three genes with hyper-methylation and reduced expression (*yaf2, casp3a, ddx56*). For gene body methylation, 42 (acute stress) and 63 (chronic stress) genes met the criteria for targeted analysis. Of these in acutely stressed fish we identified three hypo-methylated genes with respect to the control group (*cer, chpf, ahnakl*), and in chronically stressed fish we identified two hypo-methylated genes with respect to the control group (*nocta* and E3 ubiquitin-protein ligase KEG-like).

## Discussion

Our study indicates that acute and chronic environmental stressors experienced during early development have distinct effects on the gill transcriptome, methylome and on immune function in Atlantic salmon fry. We found that while acute stress had limited long-term effects on the gill transcriptome, chronic stress was associated with lasting transcriptional changes. However, both acute and chronic stress caused lasting, and contrasting, changes in the gill methylome. Crucially, early-life stress altered transcriptional response to a model pathogen challenge in a stress-specific way, with acute and chronic stress enhancing and suppressing inflammatory immune response, respectively. Our results also suggest that epigenetic changes may contribute to these modulatory effects of early life stress on immuno-competence. We identified a small proportion of genes for which an association could be made between stress-induced changes in promoter or gene body methylation and changes in expression, suggestive of a direct regulatory relationship. Furthermore, gene enrichment analysis revealed broader stress-induced epigenetic modifications within critical cellular signalling pathways involved in immune response.

### Lasting effects of stress on both the transcriptome and epigenome

Acute stress during late embryogenesis caused a lasting significant difference in the expression of fewer than 20 genes in the gill of salmon fry four months later. Previous studies have reported pronounced changes in transcription occurring immediately after similar acute temperature challenges in fish embryos and in hatched fry (Donaldson et al. 2008; Moghadam et al. 2017). However, given that the acute stress was applied four months earlier and that we observed no effects on survival and growth, the direct transcriptional response to acute, sub-lethal stress appears to be short-lived. This is consistent with known physiological and transcriptional recovery over time following acute thermal shock and other stressors (Ankley & Villeneuve 2015; Donaldson et al. 2008). In contrast, over 200 genes were differentially expressed in response to chronic stress and, functionally, these changes suggested an apparent up-regulation of active protein synthesis. Although immediate stress responses have often been associated with reduced protein synthesis in fish and mammals (Moghadam et al. 2017; Patel et al. 2002; Uren Webster & Santos 2015), cellular stress response is extremely complex and it is possible that these transcriptional changes represent a compensatory response to chronic stress. There was also evidence of up-regulation of energy metabolism, muscle differentiation and insulin-like growth factor signalling, which may be associated with a compensatory increase in growth rate in this group after the initial weight loss observed.

With regards to the epigenome, both acute and chronic stress during early life induced significant changes in gill DNA methylation profiles compared to controls. It is clear that the two different stressors induced distinct and specific alterations in the methylation profile of individual CpGs. Chronic stress induced a greater epigenetic change relative to the controls, but acute stress also caused distinct and lasting effects on the methylome, even in the absence of lasting transcriptional effects in this group. Stress has been shown to cause long-lasting alterations in methylation profiles throughout an individual’s lifetime that appear to depend on the intensity and timing of the stressor (Cao-Lei et al. 2017; Moghadam et al. 2017; Vaiserman 2015). However, to our knowledge no study has examined the contrasting effects of acute vs chronic stress on the fish transcriptome and epigenome. While the acute and chronic stress resulted in quite distinct methylation patterns of individual CpGs, functional analysis of the genes which contained or neighboured DMCpGs revealed enrichment of very similar cellular processes for both types of stress. A large number of terms related to cellular signalling pathways and their regulation were enriched for both stress groups, particularly glutamate, calcium and Rho-GTPase signalling. Epigenetic modification appears to explain the dysregulation of the neurotransmitter glutamate, commonly observed with stress-induced disorders in the mammalian brain (Nasca et al. 2015b). Glutamate also has an important signalling role in peripheral tissues, including the fish gill (Sundin et al. 2003) and it is possible that it might represent an important, wider target of epigenetic regulation. Cellular adhesion was also one of the most enriched terms in both stress groups, reflecting differential methylation of CpGs in a large number of genes encoding cadherins and protocadherins, as well as integrins, laminins and fibronectin. Cellular adhesion is critical for signal transduction as well as maintaining structure in multicellular tissues, and altered epigenetic regulation of these components has been reported for several autoimmune disorders and cancers (Zhang & Zhang 2015). This differential methylation of similar signalling pathways by both acute and chronic early life stress suggests that intra- and inter-cellular signal transduction may be a common target of stress-induced epigenetic regulation, with the potential to influence an extremely diverse array of cellular processes. However, it seems likely that the precise location and nature of CpG methylation change, which was distinct between acute and chronic stress, accounts for the fine tuning of epigenetic regulation and the resultant specific effects on transcription.

### Stress-induced changes in immune response to a model pathogen exposure

There was a very pronounced transcriptional effect of LPS in the gill in all exposed fish across treatment groups, characterising an extensive inflammatory response. Pro-inflammatory cytokines (TNF, IFN_γ_, TGFβ, IL-1b), which are typical markers of LPS regulation (Frost et al. 2002), together with many other cytokines, their regulatory factors and receptors were differentially expressed. Several pathogen-associated molecular pattern (PAMP) recognition signalling pathways (NOD-like receptor, RIG-I-like receptor and Toll-like receptor signalling) were also functionally enriched, although there is no known specific recognition receptor for LPS in fish (Sepulcre et al. 2009). Mucus provides a vital first line of pathogen defence (Linden et al. 2008), and a number of mucins (muc-2, 5, 7, 12, 17 and 19) were amongst the most up-regulated genes, together with other mucus components essential for immune response including antimicrobial factors (hepcidin, cathepsin, cathelicidin), lectins and complement factors. Transcriptional regulation and cellular signalling pathways were strongly enriched amongst genes up-regulated by LPS exposure, reflecting the diversity and complexity of the immune response. In particular, protein tyrosine kinases and phosphatases that are key regulators of signal transduction cascades (Lemmon & Schlessinger 2010) were extensively up-regulated. Furthermore, processes associated with the extracellular matrix (ECM), which provides cellular support and facilitates cellular signalling and adhesion (Bonnans et al. 2014; Naba et al. 2016; Sorokin 2010), were strongly enriched. Remodelling of the ECM is stimulated by inflammatory cytokines and has a crucial role in inflammatory response, through immune cell recruitment, activation and proliferation (Sorokin 2010). A large number of genes encoding structural components of the ECM, including many collagens, elastins, integrins, cadherins, laminins, thromobospondins and fibronectins, were up-regulated by LPS. We also found enrichment of protein transport and exocytosis, which are involved in secretion of these ECM components, and seriene/threonine proteases and matrix metallopeptidases which are important for their cleavage and activation (Naba et al. 2016). In contrast, processes related to cell division (cell cycle, DNA replication and repair) as well as protein, lipid and energy metabolism, were most strongly enriched amongst down-regulated genes. It seems likely that compensatory suppression of these processes, which are critical for the normal maintenance of tissue function and order (Ragkousi & Gibson 2014), facilitate the pronounced, acute immune response observed (Kan & Hodgkin 2014).

Transcriptional response to LPS exposure in the control group characterised a typical inflammatory immune response. We identified a significant interaction between stress and LPS response, for both the acute and chronic stressors. For fish exposed to acute cold shock during late embryogenesis, transcriptional response to LPS was of greater magnitude, but functionally similar to that of the control group. The vast majority of LPS-responsive genes identified in control fish were also differentially expressed in acutely stressed fish, and a considerable number of these were regulated to a significantly greater extent. Additional responsive genes included a number of other cytokines and their receptors, mucins and ECM components. Processes related to lipid biosynthesis were particularly over-represented amongst additional LPS-responsive genes in acutely stressed fish, including genes involved in *srebp1* signalling and sphingolipid metabolism, both of which are critical in mediating the membrane-dependent receptor based regulation of the innate immune response (Köberlin et al. 2015). This suggests that acute stress during late embryogenesis enhanced subsequent immune response, while in contrast chronic stress appeared to depress the transcriptional response to LPS, as more than 200 genes were significantly less responsive to the pathogen challenge than in the control group. These included a number of the typical pro-inflammatory response markers and many genes involved in processes such as signal transduction and ECM reorganisation, which were identified as central to the main LPS response in control fish. These results are consistent with previous reports of enhancing and suppressive effects of acute and chronic stress respectively on immuno-competence in mammals and fish (Dhabhar 2014; Glaser & Kiecolt-Glaser 2005; Smith et al. 2016; Tort 2011). While chronic stress is widely known to impair immune function, by altering the balance and activity of immune cells and cytokines, acute physiological stress can have an adaptive role, preparing organisms to deal with subsequent challenges (Monaghan & Haussmann 2015). It is thought that mild, acute stress can enhance both innate and adaptive immunity, by increasing the production and maturation of immune cells and cytokines, especially when applied during key periods of immune activation (Dhabhar 2014; Glaser & Kiecolt-Glaser 2005; Tort 2011).

### An epigenetic basis for lasting stress effects?

Epigenetic mechanisms are known to mediate lasting effects of early life stress on physiology, behaviour and disease outcomes in mammalian models on a gene-specific basis (Cao-Lei et al. 2017; Vaiserman 2015). For example, reduced methylation in the promoter of the glucocorticoid receptor, *Nr3c1,* due to early life stress is known to cause an increase in its expression in the brain, with lasting physiological and behavioural effects (Turecki & Meaney 2016). However, interpreting genome-wide associative patterns between DNA methylation and gene-expression is challenging due to the complexity of the different layers of epigenetic regulation (Hunter et al. 2015). Evidence suggests that the relationship between DNA methylation and gene expression varies widely across the genome, and occurs on a gene-specific basis (Hu et al. 2016; Jones 2012). Here we found evidence of a significant, transcriptome-wide negative correlation between DNA methylation level in putative promoters and gene expression, which is consistent with previous reports in mammals and fish (Huang et al. 2014; Wang et al. 2017; Zhong et al. 2016). In contrast, there was no linear relationship but some evidence of a more complex relationship between gene-body methylation and gene expression, which again is consistent with previous reports for mammals and plants, where non-monotonic relationships between gene methylation and expression have been reported (Jjingo et al. 2012; Lim et al. 2017; Wang et al. 2015; Zemach et al. 2010). However, transcriptome-wide, the relationship between gene expression and DNA methylation was very variable among individual genes.

Given the marked effects of both acute and chronic early life stress on the gill methylome, we hypothesised that stress-induced changes in DNA methylation of putative promoter regions and/or gene bodies could influence baseline transcription and also the rapid transcriptional response to a pathogenic challenge. We identified a small proportion of genes for which there was an association between stress-induced changes in baseline DNA methylation and transcription. These included 20 different lncRNAs, perhaps reflecting the complex and interactive nature of epigenetic modifications, since lncRNAs constitute an additional layer of epigenetic regulation at the transcriptional and post-transcriptional level (Zhao et al. 2016). There were also 12 genes which were similarly influenced by both types of stress, and overall functional analysis again revealed enrichment of ion transport and cellular signalling pathways. Similarly, we found evidence of stress-induced methylation differences (promoters or gene bodies) for a small proportion of the genes for which a significant interaction between stress and LPS response was identified. These included a number of genes involved in ubiquitination, which regulates a wide range of biological processes including the immune system, and transcriptional regulation.

Our results suggest that direct associations between promoter or gene body methylation and expression are likely to occur on a gene-specific basis. This small proportion of genes consistently appears to include components of key signalling and regulatory pathways, with the potential therefore to influence a diverse array of cellular processes. We found limited direct evidence for stress-induced alterations in the methylome corresponding to observed alterations in immune response to LPS. However, there was a marked functional overlap between gene pathways with stress-induced changes in methylation and those central to the inflammatory immune response to LPS, namely terms related to cellular adhesion/the ECM and signal transduction. This suggests that less direct mechanisms of epigenetic regulation involving DNA methylation may play a wider role in mediating the long-lasting effects of both acute and chronic stress on the immune response to pathogen challenge. These mechanisms may include DNA methylation in other features such as lncRNAs and far-distant enhancer regions, which have variable and context-specific regulatory effects on gene expression, as well as interactive effects between DNA methylation and other epigenetic modifications.

### Conclusions and potential implications

In summary, we found that acute and chronic stress during early life induced contrasting effects on gill transcription and immune response in Atlantic salmon. Acute and chronic stress also induced considerable changes in the baseline methylome, including modulation of similar cellular signalling pathways suggesting that these may be common targets of stress-induced epigenetic regulation with the potential for far-reaching effects on cellular processes. However, the specific patterns of methylation change at the individual CpG level were very different between acute and chronically stressed fish, suggesting distinction in fine level epigenetic regulation. As expected, we found that stress-induced changes in the methylome were only directly associated with transcriptional differences, and transcriptional responses to LPS, for a small proportion of genes. However, at the gene-pathway level, we present evidence for stress-induced differential methylation in the key signalling and regulatory networks involved in transcriptional response to a pathogen challenge. This suggests that stress may influence the immune response through wider, less direct, epigenetic mechanisms.

These results have important implications for health and disease management of farmed fish populations, which are commonly exposed to multiple stressors and infection challenges. They highlight the importance of considering the long-lasting effects of early life stress, even when no obvious effects on growth or body condition are apparent, and suggest that early-life stress has considerable effects on immuno-competence and disease susceptibility. Such knowledge could be used to harness the potentially stimulatory effects of acute stress on the immune system of Atlantic salmon and other commercially important fish. Our study provides the first evidence that direct and indirect epigenetic mechanisms may play a role in mediating the lasting effects of early-life stress on fish immune function.

## Materials and methods

### Ethics statement

All experiments were performed with the approval of the Swansea Animal Welfare and Ethical Review Body (AWERB; approval number IP-1415-6) and infection challenges were approved by Cardiff University Animal Ethics Committee and conducted under a UK Home Office License (PPL 302876).

### Stress experiments

Atlantic salmon eggs were assigned at random to three experimental treatments: control, acute environmental stress and chronic environmental stress, with two replicate groups of 500 eggs per treatment. To rule out potential family effects, eggs were obtained from 10 different families (1:1 crosses) and these were equally assigned to each experimental group. We used an acute stressor which has previously been shown to induce repeatable effects on the teleost transcriptome and methylome, and is relevant to temperature-shocking procedures commonly used in aquaculture to screen eggs (Moghadam et al. 2017). During late embryogenesis (360 DD), embryos were immersed in iced water (0.2 °C) for five minutes and then exposed to air (12 °C) for five minutes before being returned to normal water temperature (9 °C). For the chronic stress group, larvae were reared in fry troughs without the artificial hatching substrate (Astroturf) used in the control and acute stress groups for the duration of the experiment (four months). Artificial substrate is routinely used in salmon farming to mimic the natural substrate, and provides support and shelter to fish larvae. Salmonid larvae reared in bare troughs tend to show elevated cortisol levels, developmental abnormalities and impaired growth (Bates et al. 2014; Hansen 1985). Full details on fish husbandry are given in the supporting information.

Daily mortalities of embryos, larvae and fry were recorded, and growth was monitored based on a subset of 20 euthanised individuals from each of the six replicate troughs at four time-points; 492, 748, 1019 and 1323 DD. At the final sampling point (1532 DD), mass and fork length of 20 fish from each tank were determined and used to calculate Fulton’s condition factor (Froese 2006). All gill arches from both sides of each fish were dissected out and stored in RNAlater (Sigma Aldrich, UK) at 4°C for 24 h followed by longer term storage at −20 °C for subsequent RNA/DNA extraction. At each of the five sampling points separately, the effects of stress treatment (control, acute, chronic) on fish mass, as well as condition factor at the final sampling point, was assessed using linear mixed effect models (lme function in nmle (Pinheiro et al. 2017)) in R version 3.3.3 (R_Core_Team 2014) using tank identity as a random factor to account for variation between replicate tanks.

### Immuno-stimulation experiment

To assess the effect of acute and chronic environmental stress on immune response, a subset of salmon fry from each stress group and the control group were exposed to lipopolysaccharide (LPS), a pathogen-associated molecular pattern, mimicking a bacterial infection, at the end of the experimental period (1532 DD). Six fry from each replicate tank (12 per group) were exposed to 20 μg/ml LPS obtained from *Pseudomonas aeruginosa* (Sigma Aldrich, UK) for 24 h in 0.5 L tanks, each containing a static volume of aerated water (Sundaram et al. 2012). Fish were visually monitored during the course of the experiment and any adverse effects on behaviour and physiology recorded. After 24h of exposure, fry were euthanised, weighed and measured, and all gill arches were dissected out and stored in RNAlater. Gills were selected as a target tissue for analysis because they have a critical role in immune defence, and are a primary target of waterborne pathogens (Uribe et al. 2011).

### Transcriptome and methylome sequencing

Matched transcriptome and methylome analysis of the gill was performed at the final sampling point (1532 DD) for a total of eight fish in each of the two stress groups and the control group (24 fish in total, including four from each replicate tank). Transcriptome analysis was also performed on the gills of eight LPS-exposed fish from each of the three experimental groups (24 fish; four per replicate tank). RNA and DNA were simultaneously extracted using the Qiagen AllPrep DNA/RNA Mini Kit, and all libraries were prepared using high quality RNA and DNA (full details given in supporting information). Transcriptomic analysis was conducted using RNA-seq; the 48 libraries were prepared using the Illumina TruSeq RNA preparation kit and sequenced on an Illumina NextSeq500 platform (76bp paired-end reads). Methylation analysis was performed using Reduced Representation Bisulfide Sequencing (RRBS); the 24 libraries were prepared using the Diagenode Premium RRBS Kit, and sequenced on Illumina NextSeq 500 (76 bp single-end reads).

### Bioinformatics analysis

All raw sequence reads have been deposited to the European Nucleotide Archive under the accession numbers PRJEB25636 and PRJEB25637, and full details of bioinformatics analyses performed are provided in the supporting information.

Briefly, for the transcriptomics analysis, following quality screening and filtering using Trimmomatic (Bolger et al. 2014), high quality reads were then aligned to the Atlantic salmon genome (v GCF_000233375.1_ICSAG_v2; (Davidson et al. 2010; Lien et al. 2016)) using HISAT2 (v 2.1.0; (Kim et al. 2015)), followed by transcript reconstruction and assembly using StringTie (v1.3.3) (Pertea et al. 2015) and extraction of non-normalised transcript read counts. Differentially expressed genes in response to stress and LPS exposure were identified using a multifactorial design in DeSeq2 (Love et al. 2014), including the main effects of stress and LPS exposure, and their interaction, and accounting for potential variation between replicate tanks. Genes were considered significantly differentially expressed at FDR <0.05. Hierarchical clustering of all genes significantly regulated by LPS, and all genes for which a significant interaction between stress and LPS response was identified, was performed using an Euclidean distance metric and visualised using the Pheatmap package in R (Kolde 2015). Functional enrichment analysis of differentially regulated genes was performed using DAVID (v 6.8; (Huang et al. 2008)), using zebrafish orthologs for improved functional annotation.

For the methylation analysis, initial read quality filtering was performed using TrimGalore (Kreger 2016) before high quality reads were aligned to the Atlantic salmon reference genome and cytosine methylation calls extracted using Bismark v 0.17.0 (Krueger & Andrews 2011). Mapped data were then processed using SeqMonk (Andrews 2007), considering only methylation within CpG context, and only including CpGs with a minimum coverage of 10 reads in each of the 24 samples in the analysis. Differentially methylated CpGs (DMCpGs) were identified using logistic regression (FDR<0.01 and >20% minimal CpG methylation difference (|ΔM|)). For each DMCpG, we identified the genomic location (gene body, promoter region (≤1500 bp upstream of the transcription start site (TSS)), or intergenic region) and the context location (CpG island (≥200 bp with GC % ≥ 55% and an observed-to-expected CpG ratio of ≥ 65%), CpG shore (up to 2 kb of a CpG island), CpG shelf (up to 2 kb of a CpG shore)). For the DMCpGs that were within a gene, or within 2 kb (upstream or downstream) of the TSS or transcription termination site (TTS) respectively, we also performed gene function enrichment analysis as described above. To generate a more stringent list of DMCpGs for further cluster analysis between stress groups, we additionally ran t-tests for each paired comparison using a threshold of p<0.01, to identify DMCpGs shared by both statistical methods.

### Transcriptome-methylome integration

To explore the relationship between the methylome and the transcriptome we performed targeted DNA methylation analysis for putative gene regulatory regions and for gene bodies, and in each case investigated the relationship between total methylation level and gene expression. For the analysis of gene bodies, we only used gene bodies containing ≥ 5 CpGs, each with ≥ 10 reads per CpG, in all 24 samples (10,017 genes covered out of 61,274 overall expressed genes; 16.3%). To increase the number of p.promoter regions included in the analysis, we used a lower threshold (regions containing ≥ 3 CpGs, each with ≥ 5 reads per CpG, in all 24 samples; 5,422 gene promoter regions covered; 8.8% expressed genes).

We performed a Spearman correlation between mean gene expression and mean DNA methylation within p.promoters and, separately, within gene bodies, for all covered genes in the control group. We also aimed to identify genes for which early life stress influenced both DNA methylation and gene transcription. For all expressed genes at the baseline time-point (i.e. not exposed to LPS), we plotted gene expression difference in each of the stress groups relative to the control group (delta expression) against the respective difference in gene body methylation and in p.promoter methylation (delta methylation). We identified genes with a marked effect of stress on both expression and methylation (>2 fold delta expression and >5% difference in methylation), based on previously described thresholds (e.g. Lussier et al. 2018; Muurinen et al. 2017) and performed functional enrichment analysis as before. We also performed a Chi-square test incorporating Yates’ correction to test distribution of hypo-methylated/up-regulated, hyper-methylated/up-regulated, hypo-methylated/down-regulated and hyper-methylated/down-regulated genes.

We also aimed to investigate the possible role of stress-induced changes in DNA methylation in influencing transcriptional response to the immune challenge (LPS). Therefore, for all genes for which a significant interaction between stress and transcriptional response to LPS exposure was identified, we plotted delta expression following exposure to LPS relative to that in the control group against delta baseline methylation relative to the control group, and identified genes with a >5% difference in baseline methylation.

## Acknowledgements

We are grateful to Sam Fieldwick for assistance with sampling and Dr Angela Marchbank for facilitating the RRBS sequencing. This work was funded by a BBSRC-NERC Aquaculture grant (BB/M026469/1) and by the Welsh Government and Higher Education Funding Council for Wales (HEFCW) through the Sêr Cymru National Research Network for Low Carbon Energy and Environment (NRN-LCEE).

## Author contributions

CGL, SC, TUW, SM, CvO and JC designed the study; AH provided materials for the experiment; TUW and DRB collected and analysed the data with assistance from SM and CvO; TUW, DRB, CGL and SC wrote the manuscript. All authors contributed to the final version of the manuscript.

## References

Andrews, S. 2007 SeqMonk: A tool to visualise and analyse high throughput mapped sequence data. https://www.bioinformatics.babraham.ac.uk/projects/seqmonk/.

Ankley, G. T. & Villeneuve, D. L. 2015 Temporal Changes in Biological Responses and Uncertainty in Assessing Risks of Endocrine-Disrupting Chemicals: Insights from Intensive Time-Course Studies with Fish. Toxicological Sciences 144, 259–275.

Baerwald, M. R., Meek, M. H., Stephens, M. R., Nagarajan, R. P., Goodbla, A. M., Tomalty, K. M. H., Thorgaard, G. H., May, B. & Nichols, K. M. 2016 Migration-related phenotypic divergence is associated with epigenetic modifications in rainbow trout. Molecular Ecology 25, 1785–1800.

Bates, L., Boucher, M. & Shrimpton, J. 2014 Effect of temperature and substrate on whole body cortisol and size of larval White Sturgeon (*Acipenser transmontanus* Richardson, 1836). Journal of Applied Ichthyology 30, 1259–1263.

Bolger, A. M., Lohse, M. & Usadel, B. 2014 Trimmomatic: a flexible trimmer for Illumina sequence data. Bioinformatics 30, 2114–20.

Bonnans, C., Chou, J. & Werb, Z. 2014 Remodelling the extracellular matrix in development and disease. Nature Reviews Molecular Cell Biology 15, 786–801.

Calcagni, E. & Elenkov, I. 2006 Stress system activity, innate and T helper cytokines, and susceptibility to immune-related diseases. Annals of the New York Academy of Sciences 1069, 62–76.

Cao-Lei, L., de Rooij, S. R., King, S., Matthews, S. G., Metz, G. A. S., Roseboom, T. J. & Szyf, M. 2017 Prenatal stress and epigenetics. Neuroscience & Biobehavioral Reviews Epub ahead of print.

Castro, R., Jouneau, L., Tacchi, L., Macqueen, D. J., Alzaid, A., Secombes, C. J., Martin, S. A. M. & Boudinot, P. 2015 Disparate developmental patterns of immune responses to bacterial and viral infections in fish. Scientific Reports 5, 15458.

Chatterjee, A., Ozaki, Y., Stockwell, P. A., Horsfield, J. A., Morison, I. M. & Nakagawa, S. 2013 Mapping the zebrafish brain methylome using reduced representation bisulfite sequencing. Epigenetics 8, 979–89.

Chrousos, G. P. 2009 Stress and disorders of the stress system. Nature Reviews Endocrinology 5, 374–382.

Cruceanu, C., Matosin, N. & Binder, E. B. 2017 Interactions of early-life stress with the genome and epigenome: from prenatal stress to psychiatric disorders. Current Opinion in Behavioral Sciences 14, 167–171.

Davidson, W. S., Koop, B. F., Jones, S. J., Iturra, P., Vidal, R., Maass, A., Jonassen, I., Lien, S. & Omholt, S. W. 2010 Sequencing the genome of the Atlantic salmon *(Salmo salar*). Genome Biology 11, 403–410.

Dhabhar, F. S. 2009 Enhancing versus Suppressive Effects of Stress on Immune Function: Implications for Immunoprotection and Immunopathology. Neuroimmunomodulation 16, 300–317.

Dhabhar, F. S. 2014 Effects of stress on immune function: the good, the bad, and the beautiful. Immunologic Research 58, 193–210.

Donaldson, M., Cooke, S., Patterson, D. & Macdonald, J. 2008 Cold shock and fish. Journal of Fish Biology 73, 1491–1530.

Elliott, J. 1989 The critical-period concept for juvenile survival and its relevance for population regulation in young sea trout, *Salmo trutta*. Journal of Fish Biology 35, 91– 98.

Froese, R. 2006 Cube law, condition factor and weight–length relationships: history, meta-analysis and recommendations. Journal of Applied Ichthyology 22, 241–253.

Frost, R. A., Nystrom, G. J. & Lang, C. H. 2002 Lipopolysaccharide regulates proinflammatory cytokine expression in mouse myoblasts and skeletal muscle. American Journal of Physiology-Regulatory, Integrative and Comparative Physiology 283, R698–R709.

Glaser, R. & Kiecolt-Glaser, J. K. 2005 Stress-induced immune dysfunction: implications for health. Nature Reviews Immunology 5, 243–251.

Hansen, T. 1985 Artificial hatching substrate: Effect on yolk absorption, mortality and growth during first feeding of sea trout (*Salmo trutta*). Aquaculture 46, 275–285.

Hu, Y., Huang, K., An, Q., Du, G., Hu, G., Xue, J., Zhu, X., Wang, C.-Y., Xue, Z. & Fan, G. 2016 Simultaneous profiling of transcriptome and DNA methylome from a single cell. Genome Biology 17, 88–94.

Huang, D. W., Sherman, B. T. & Lempicki, R. A. 2008 Systematic and integrative analysis of large gene lists using DAVID bioinformatics resources. Nature Protocols 4, 44–57.

Huang, Y.-Z., Sun, J.-J., Zhang, L.-Z., Li, C.-J., Womack, J. E., Li, Z.-J., Lan, X.-Y., Lei, C.-Z., Zhang, C.-L., Zhao, X. & Chen, H. 2014 Genome-wide DNA Methylation Profiles and Their Relationships with mRNA and the microRNA Transcriptome in Bovine Muscle Tissue (*Bos taurine*). Scientific Reports 4, 6546–6552.

Hunter, R. G., Gagnidze, K., McEwen, B. S. & Pfaff, D. W. 2015 Stress and the dynamic genome: Steroids, epigenetics, and the transposome. Proceedings of the National Academy of Sciences 112, 6828–6833.

Jjingo, D., Conley, A. B., Yi, S. V., Lunyak, V. V. & Jordan, I. K. 2012 On the presence and role of human gene-body DNA methylation. Oncotarget 3, 462–474.

Jones, P. A. 2012 Functions of DNA methylation: islands, start sites, gene bodies and beyond. Nature Reviews Genetics 13, 484–92.

Kan, A. & Hodgkin, P. D. 2014 Mechanisms of cell division as regulators of acute immune response. Systems and Synthetic Biology 8, 215–221.

Kim, D., Langmead, B. & Salzberg, S. L. 2015 HISAT: a fast spliced aligner with low memory requirements. Nature Methods 12, 357–360.

Köberlin, Marielle S., Snijder, B., Heinz, Leonhard X., Baumann, Christoph L., Fauster, A., Vladimer, Gregory I., Gavin, A.-C. & Superti-Furga, G. 2015 A Conserved Circular Network of Coregulated Lipids Modulates Innate Immune Responses. Cell 162, 170– 183.

Kolde, R. 2015 Pretty Heatmaps. R package version 3.1-131.

Kreger, F. 2016 TrimGalore. A wrapper around Cutadapt and FastQC to consistently apply adapter and quality trimming to FastQ files, with extra functionality for RRBS data. https://www.bioinformatics.babraham.ac.uk/projects/trim_galore/.

Krueger, F. & Andrews, S. R. 2011 Bismark: a flexible aligner and methylation caller for Bisulfite-Seq applications. Bioinformatics 27, 1571–1572.

Lemmon, M. A. & Schlessinger, J. 2010 Cell signaling by receptor tyrosine kinases. Cell 141, 1117–1134.

Lien, S., Koop, B. F., Sandve, S. R., Miller, J. R., Kent, M. P., Nome, T., Hvidsten, T. R., Leong, J. S., Minkley, D. R., Zimin, A., Grammes, F., Grove, H., Gjuvsland, A., Walenz, B., Hermansen, R. A., von Schalburg, K., Rondeau, E. B., Di Genova, A., Samy, J. K. A., Olav Vik, J., Vigeland, M. D., Caler, L., Grimholt, U., Jentoft, S., Inge Våge, D., de Jong, P., Moen, T., Baranski, M., Palti, Y., Smith, D. R., Yorke, J. A., Nederbragt, A. J., Tooming-Klunderud, A., Jakobsen, K. S., Jiang, X., Fan, D., Hu, Y., Liberles, D. A., Vidal, R., Iturra, P., Jones, S. J. M., Jonassen, I., Maass, A., Omholt, S. W. & Davidson, W. S. 2016 The Atlantic salmon genome provides insights into rediploidization. Nature 533, 200–205.

Lim, Y. C., Li, J., Ni, Y., Liang, Q., Zhang, J., Yeo, G. S., Lyu, J., Jin, S. & Ding, C. 2017 A complex association between DNA methylation and gene expression in human placenta at first and third trimesters. PloS One 12, e0181155.

Linden, S. K., Sutton, P., Karlsson, N. G., Korolik, V. & McGuckin, M. A. 2008 Mucins in the mucosal barrier to infection. Mucosal Immunology 1, 183–190.

Love, M. I., Huber, W. & Anders, S. 2014 Moderated estimation of fold change and dispersion for RNA-seq data with DESeq2. Genome Biology 15, 550–555.

Lussier, A. A., Morin, A. M., MacIsaac, J. L., Salmon, J., Weinberg, J., Reynolds, J. N., Pavlidis, P., Chudley, A. E. & Kobor, M. S. 2018 DNA methylation as a predictor of fetal alcohol spectrum disorder. Clinical Epigenetics 10, 5–10.

Mattson, M. P. & Calabrese, E. J. 2009 Hormesis: a revolution in biology, toxicology and medicine: Springer Science & Business Media.

McGowan, P. O., Sasaki, A., D’alessio, A. C., Dymov, S., Labonté, B., Szyf, M., Turecki, G. & Meaney, M. J. 2009 Epigenetic regulation of the glucocorticoid receptor in human brain associates with childhood abuse. Nature Neuroscience 12, 342–348.

Moghadam, H. K., Johnsen, H., Robinson, N., Andersen, Ø., H. Jørgensen, E., Johnsen, H. K., Bæhr, V. J. & Tveiten, H. 2017 Impacts of Early Life Stress on the Methylome and Transcriptome of Atlantic Salmon. Scientific Reports 7, 5023.

Monaghan, P. & Haussmann, M. F. 2015 The positive and negative consequences of stressors during early life. Early Human Development 91, 643–647.

Muurinen, M., Hannula-Jouppi, K., Reinius, L. E., Söderhäll, C., Merid, S. K., Bergström, A., Melén, E., Pershagen, G., Lipsanen-Nyman, M., Greco, D. & Kere, J. 2017 Hypomethylation of HOXA4 promoter is common in Silver-Russell syndrome and growth restriction and associates with stature in healthy children. Scientific Reports 7, 15693–99.

Naba, A., Clauser, K. R., Ding, H., Whittaker, C. A., Carr, S. A. & Hynes, R. O. 2016 The extracellular matrix: Tools and insights for the “omics” era. Matrix Biology 49, 10–24.

Nasca, C., Bigio, B., Zelli, D., Nicoletti, F. & McEwen, B. S. 2015a Mind the gap: glucocorticoids modulate hippocampal glutamate tone underlying individual differences in stress susceptibility. Molecular Psychiatry 20, 755–763.

Nasca, C., Zelli, D., Bigio, B., Piccinin, S., Scaccianoce, S., Nisticò, R. & McEwen, B. S. 2015b Stress dynamically regulates behavior and glutamatergic gene expression in hippocampus by opening a window of epigenetic plasticity. Proceedings of the National Academy of Sciences 112, 14960–14965.

Papadopoulou, A., Siamatras, T., Delgado-Morales, R., Amin, N. D., Shukla, V., Zheng, Y. L., Pant, H. C., Almeida, O. F. X. & Kino, T. 2015 Acute and chronic stress differentially regulate cyclin-dependent kinase 5 in mouse brain: implications to glucocorticoid actions and major depression. Translational Psychiatry 5, e578.

Patel, J., McLeod, L. E., Vries, R. G., Flynn, A., Wang, X. & Proud, C. G. 2002 Cellular stresses profoundly inhibit protein synthesis and modulate the states of phosphorylation of multiple translation factors. European Journal of Biochemistry 269, 3076–85.

Pertea, M., Pertea, G. M., Antonescu, C. M., Chang, T.-C., Mendell, J. T. & Salzberg, S. L. 2015 StringTie enables improved reconstruction of a transcriptome from RNA-seq reads. Nature Biotechnology 33, 290–295.

Pinheiro, J., Bates, D., DebRoy, S., Sarker, D. & Team, R. C. 2017 nlme: Linear and Nonlinear Mixed Effects Models: R package version 3.1-131.

R_Core_Team. 2014 R: A language and environment for statistical computing. Vienna, Austria.: R Foundation for Statistical Computing.

Ragkousi, K. & Gibson, M. C. 2014 Cell division and the maintenance of epithelial order. The Journal of Cell Biology 207, 181–188.

Schreck, C. B., Tort, L., Farrell, A. P. & Brauner, C. J. 2016 Biology of Stress in Fish: Academic Press.

Sepulcre, M. P., Alcaraz-Pérez, F., López-Muñoz, A., Roca, F. J., Meseguer, J., Cayuela, M. L. & Mulero, V. 2009 Evolution of lipopolysaccharide (LPS) recognition and signaling: fish TLR4 does not recognize LPS and negatively regulates NF-κB activation. The Journal of Immunology 182, 1836–1845.

Silberman, D. M., Acosta, G. B. & Zorrilla Zubilete, M. A. 2016 Long-term effects of early life stress exposure: Role of epigenetic mechanisms. Pharmacological Research 109, 64–73.

Smith, B. L., Schmeltzer, S. N., Packard, B. A., Sah, R. & Herman, J. P. 2016 Divergent effects of repeated restraint versus chronic variable stress on prefrontal cortical immune status after LPS injection. Brain, Behavior, and Immunity 57, 263–270.

Sorokin, L. 2010 The impact of the extracellular matrix on inflammation. Nature Reviews Immunology 10, 712–723.

Stentiford, G. D., Sritunyalucksana, K., Flegel, T. W., Williams, B. A., Withyachumnarnkul, B., Itsathitphaisarn, O. & Bass, D. 2017 New Paradigms to Help Solve the Global Aquaculture Disease Crisis. PLoS pathogens 13, e1006160.

Sundaram, A. Y., Consuegra, S., Kiron, V. & Fernandes, J. M. 2012 Positive selection pressure within teleost Toll-like receptors tlr21 and tlr22 subfamilies and their response to temperature stress and microbial components in zebrafish. Molecular Biology Reports 39, 8965–75.

Sundin, L., Turesson, J. & Taylor, E. W. 2003 Evidence for glutamatergic mechanisms in the vagal sensory pathway initiating cardiorespiratory reflexes in the shorthorn sculpin *Myoxocephalus scorpius*. Journal of Experimental Biology 206, 867–876.

Tort, L. 2011 Stress and immune modulation in fish. Developmental & Comparative Immunology 35, 1366–1375.

Turecki, G. & Meaney, M. J. 2016 Effects of the Social Environment and Stress on Glucocorticoid Receptor Gene Methylation: A Systematic Review. Biological Psychiatry 79, 87–96.

Uren Webster, T. M. & Santos, E. M. 2015 Global transcriptomic profiling demonstrates induction of oxidative stress and of compensatory cellular stress responses in brown trout exposed to glyphosate and Roundup. BMC Genomics 16, 32–38.

Uribe, C., Folch, H., Enriquez, R. & Moran, G. 2011 Innate and adaptive immunity in teleost fish: a review. Veterinarni Medicina 56, 486–503.

Vaiserman, A. M. 2015 Epigenetic programming by early-life stress: Evidence from human populations. Developmental Dynamics 244, 254–265.

Vindas, M. A., Madaro, A., Fraser, T. W. K., Höglund, E., Olsen, R. E., Øverli, Ø. & Kristiansen, T. S. 2016 Coping with a changing environment: the effects of early life stress. Royal Society Open Science 3.

Wang, H., Beyene, G., Zhai, J., Feng, S., Fahlgren, N., Taylor, N. J., Bart, R., Carrington, J. C., Jacobsen, S. E. & Ausin, I. 2015 CG gene body DNA methylation changes and evolution of duplicated genes in cassava. Proceedings of the National Academy of Sciences 112, 13729–13734.

Wang, H., Wang, J., Ning, C., Zheng, X., Fu, J., Wang, A., Zhang, Q. & Liu, J.-F. 2017 Genome-wide DNA methylation and transcriptome analyses reveal genes involved in immune responses of pig peripheral blood mononuclear cells to poly I:C. Scientific Reports 7, 9709–9715.

Zemach, A., Kim, M. Y., Silva, P., Rodrigues, J. A., Dotson, B., Brooks, M. D. & Zilberman, D. 2010 Local DNA hypomethylation activates genes in rice endosperm. Proceedings of the National Academy of Sciences 107, 18729–18734.

Zhang, Z. & Zhang, R. 2015 Epigenetics in autoimmune diseases: Pathogenesis and prospects for therapy. Autoimmunity Reviews 14, 854–63.

Zhao, Y., Sun, H. & Wang, H. 2016 Long noncoding RNAs in DNA methylation: new players stepping into the old game. Cell & Bioscience 6, 45.

Zhong, Z., Du, K., Yu, Q., Zhang, Y. E. & He, S. 2016 Divergent DNA Methylation Provides Insights into the Evolution of Duplicate Genes in Zebrafish. G3: Genes|Genomes|Genetics 6, 3581–3591.

